# Bartender: an ultrafast and accurate clustering algorithm to count barcode and amplicon reads

**DOI:** 10.1101/068916

**Authors:** Lu Zhao, Zhimin Liu, Sasha F. Levy, Song Wu

## Abstract

Barcode sequencing (bar-seq) is a high-throughput, and cost effective method to assay large numbers of lineages or genotypes in complex cell pools. Because of its advantages, applications for bar-seq are quickly growing – from using neutral random barcodes to study the evolution of microbes or cancer, to using pseudo-barcodes, such as shRNAs, sgRNAs, or transposon insertion libraries, to simultaneously screen large numbers of cell perturbations. However, the computational pipelines for bar-seq have not been well developed. Available methods, which use prior information and/or simple brute-force comparisons, are slow and often result in overclustering artifacts that group distinct barcodes together. Here, we developed Bartender: an ultrafast and accurate clustering algorithm to detect barcodes and their abundances from raw next-generation sequencing data. To improve speed and reduce unnecessary pairwise comparisons, Bartender employs a divide-and-conquer strategy that intelligently sorts barcode reads into distinct bins before performing comparisons. To improve accuracy and reduce over-clustering artifacts, Bartender employs a modified two-sample proportion test that uses information on both the cluster sequence distances and cluster sizes to make merging decisions. Additionally, Bartender includes a “multiple time point” mode, which matches barcode clusters between different clustering runs for seamless handling of time course data. For both simulated and real data, Bartender clusters millions of unique barcodes in a few minutes at high accuracy (>99.9%), and is ~100-fold faster than previous methods. Bartender is a set of simple-to-use command line tools that can be performed on a laptop.

**Availability:** Bartender is available at no charge for non-commercial use at https://github.com/LaoZZZZZ/bartender-1.1.

## Introduction

High-throughput sequencing of nucleotide barcodes (bar-seq) provides a powerful tool to assay and track dynamics of large numbers of lineages, genotypes or perturbations in complex cell pools. It generally works by growing a pool of barcoded cells under selective conditions, amplifying extracted barcodes using common primers, and sequencing barcode amplicons to quantify relative barcode frequencies in the cell pool. This approach was first applied on the *Saccharomyces cerevisiae* deletion collection, which was designed such that each individual deletion strain is marked with a unique barcode (Giaever et al., 2002; Winzeler et al., 1999). Bar-seq of these deletion strains have been used, for example, to detect gene products that are drug targets (Smith et al., 2009), and those that are important for surviving starvation (Gresham et al., 2011), and heat stress (Gibney, Lu, Caudy, Hess, & Botstein, 2013). Barcoded deletion collections have subsequently been generated in a number of bacteria and yeasts, allowing for similar massively parallel functional profiling (Han, Xu, Zhang, Peng, & Du, 2010; Hobbs, Astarita, & Storz, 2010; Noble et al., 2010; Schwarzmuller et al., 2014). Analogously, a growing number of studies in mammalian cells sequence pseudo-barcodes: short nucleotide sequences such as shRNAs or sgRNAs, that serve as both the cell-specific perturbation and the unique cell identifier for short-read sequencing (Bassik et al., 2009; Schlabach et al., 2008; Silva, Rowntree, Mekhoubad, & Lee, 2008; Sims et al., 2011; Wang, Wei, Sabatini, & Lander, 2014; Wong, Choi, Cheng, Purcell, & Lu, 2015).

In addition to the above approaches where the sequence of the barcode or pseudo-barcode is known *a priori*, more recent bar-seq studies employ barcodes of random unknown sequence. As one example, sequencing of the DNA that happens to be adjacent to a transposon in random transposon insertion libraries has been used to functionally profile gene disruptions when systematic deletion collections are unavailable (Brutinel & Gralnick, 2012; Carette et al., 2011; Gawronski, Wong, Giannoukos, Ward, & Akerley, 2009; van Opijnen, Bodi, & Camilli, 2009). A second example is the insertion of random barcode libraries into genomes to serve as neutral markers for lineage tracking studies over the course of development, evolution or cancer progression (Bhang et al., 2015; Blundell & Levy, 2014; Levy et al., 2015; Lu, Neff, Quake, & Weissman, 2011; Nguyen et al., 2015). In these studies, random barcodes are generated from primers with random bases, which are first inserted into vectors and then into cell genomes.

Despite the widespread use of bar-seq, computational pipelines for handling the resulting sequencing data have not been well developed. For barcodes of known sequence, the primary concern is mapping reads that may contain PCR or sequencing errors to the known barcodes. One naïve strategy would be to ignore reads that do not exactly match any putative barcode. However, given that some barcodes in the pool may be more prone to PCR or sequencing errors (Goren et al., 2010; Gundry & Vijg, 2012; Meyerhans, Vartanian, & Wain-Hobson, 1990; Schmitt et al., 2012), this strategy could introduce counting biases. A more sensible strategy currently employed is to compare each read to the set of putative barcodes by calculating the Hamming (Hamming, 1950) or Levenshtein distance (Levenshtein, 1966), thereby discovering the best match (Gresham et al., 2011). However, this strategy is extremely computationally expensive and computational cost grows at least quadratically with the number of putative barcodes. Additionally, *a priori* errors in the set of known barcodes (likely due to errors that occurred during Sanger sequencing of barcoded clones) have been found to be common (Smith et al., 2009), meaning that unexpected barcodes that are present in the pool may be missed. Therefore, an unbiased barcode detection strategy that does not depend on prior information is needed.

For random barcode libraries, an additional computational problem is discovering the true barcodes in the pool. That is, reads that identify a true barcode must be differentiated from reads that contain PCR or sequencing errors. Because sequences representing an exact match to a true barcode are likely to be sequenced at much higher frequencies than those with errors, one approach would be to ignore reads below a predefined frequency threshold and treat all other reads as true barcodes. However, in cell pools with a skewed barcode frequency distribution, less abundant true barcodes will fall below the threshold, while errors from extremely abundant barcodes rise above it. If each barcode in the pool is expected to be distant in sequence space from all other barcodes (e.g. >3 mismatches), a second approach is to cluster reads by their sequence similarities (Bhang et al., 2015; Levy et al., 2015; Lu et al., 2011). Here, reads that are within one or two mismatches of each other are grouped together, with the most abundant sequence likely to be the true barcode and others being errors. This approach, however, is currently extremely computationally expensive because each unique read must be compared against each other unique read, with the number of unique reads often exceeding 10^6^ in a typical Illumina HiSeq run. To avoid calculating all pairwise Hamming or Levenshtein distances, we have previously used a semi-pairwise BLAST strategy to cluster reads (Altschul, Gish, Miller, Myers, & Lipman, 1990; Levy et al., 2015). However, even this strategy is computationally expensive and may often falsely merge distinct barcodes that are close in sequence space (see Results).

Here, we developed Bartender, an ultrafast and unbiased clustering algorithm for identifying and counting barcodes and pseudo-barcodes from short read sequencing data. We use a divide-and-conquer strategy that first identifies high-quality seeds and then iterates through each seed to sort short reads into different bins for parallel processing. Reads within each bin are clustered using a computationally efficient greedy clustering algorithm. Instead of merging clusters solely on sequence similarity (Hamming or Levenshtein distance), Bartender uses additional information of the cluster sizes to prevent over-merging. Our algorithm includes handling of unique molecular identifiers (UMIs), which, if included in the sequencing reads, allow an investigator to detect and remove PCR duplicates (Kivioja et al., 2012; Levy et al., 2015). Additionally, we include a “multiple time point” mode, which uses cluster information from adjacent time points to minimize the impact of read errors on barcode trajectories. Compared with previous methods, we find Bartender to be orders of magnitude faster and more accurate for routine processing of barcodes and pseudo-barcodes.

## Results

### Bartender is flexible and ultrafast

Barcode or pseudo-barcode lengths typically range between 10 and 30 nucleotides. However, with the advent of highly complex long oligonucleotide libraries (LeProust et al., 2010), variable regions may sometimes exceed 100 nucleotides (Goodman, Church, & Kosuri, 2013; Kosuri et al., 2013). Current tools become cumbersome for analysis of these longer barcodes for two major reasons. First, longer barcodes will accumulate more errors and result in more unique reads that must be clustered. Second, longer barcodes will slow similarity comparisons between unique reads because more nucleotide positions must be accounted for. To address these problems, we built Bartender with the capability and speed to handle arbitrary barcode lengths.

First, we built an extractor tool that can quickly pull the variable region from FASTQ/FASTA files using a user-defined pattern and sequencing quality threshold to generate a pool of barcode reads for clustering (Methods). Synthesis of oligonucleotides with random regions often results in few variants with missing or additional random bases. To account for these errors, we allow users to assign random nucleotide regions of variable lengths during extraction. Our extractor tool also estimates the combined PCR and sequencing error rate by examining the frequency of nucleotide errors at user-defined invariant regions of the amplicons.

Second, the Bartender clustering tool utilizes a number of strategies to improve speed. The primary speedup comes from a divide-and-conquer binning strategy that greatly reduces the number of comparisons (Methods). Briefly, Bartender surveys the variable regions for nucleotide positions with the largest entropy (most variability) and generates a set of non-contiguous seeds (3-8 nucleotides) ranked by total entropy. These seeds are then used to sort unique reads into bins, with all reads within a bin having an identical seed. For the first seed, each unique read is treated as one cluster. Comparisons are then performed within each bin to merge similar clusters (the merging criteria are described later), and these merged clusters are used with the next seed for further merging. By using seeds with the largest entropy first, Bartender creates the maximum number of bins at the first clustering step (when the most clusters exist) and thereby minimizes the number of comparisons necessary. In most cases, the number of comparisons needed drops dramatically between rounds of merging. A secondary speedup comes from a greedy algorithm that prioritizes comparisons to the largest clusters within each bin. These large clusters are the most likely to be “true” barcodes and have a better chance of matching to smaller clusters that are more likely to be a sequence that contains a PCR or sequencing error. A last source of increased speed comes from the use of a computationally efficient comparison metric that is described in more detail later.

To measure the effect of these improvements, we compared Bartender clustering speed to the fastest existing clustering method, which performs comparisons between unique reads by BLAST (see Methods and (Levy et al., 2015)). We simulated 100,000 barcodes of a number of different realistic barcode lengths (26, 38, and 64 nucleotides), while keeping the number of total reads (~10M), and the combined PCR and sequencing error rate (2%) constant (see Supplementary Materials). For each simulation, barcode frequencies were chosen to follow an exponential distribution with a mean of 100. On a standard desktop computer, Bartender performed the clustering within two minutes in all cases, at least two orders of magnitude faster than BLAST (Figure 1A). One tunable parameter that affects Bartender performance is the seed length used to partition unique reads prior to pairwise comparisons (Methods). Longer seeds will reduce the pairwise comparisons (increase speed) by creating more bins, but, if error rates are high, may result in under-merging because similar clusters might never find themselves in the same bin. By testing Bartender performance across different seed lengths, we find that longer seeds do indeed increase speed, however the increments become marginal when seed length is above 5 (Figure 1B). We next performed these same speed tests on a typical laptop computer with less computing power, and found that performance is only moderately diminished (< 2-fold slower, Figure 1B). In both cases, a large portion of processing time used by Bartender is attributed to the Input/Output (I/O), particularly at longer seed lengths. Taken together, these results indicate Bartender can be used to process most realistic barcode libraries on most computers quickly, usually in just a few minutes.

**Figure 1.**
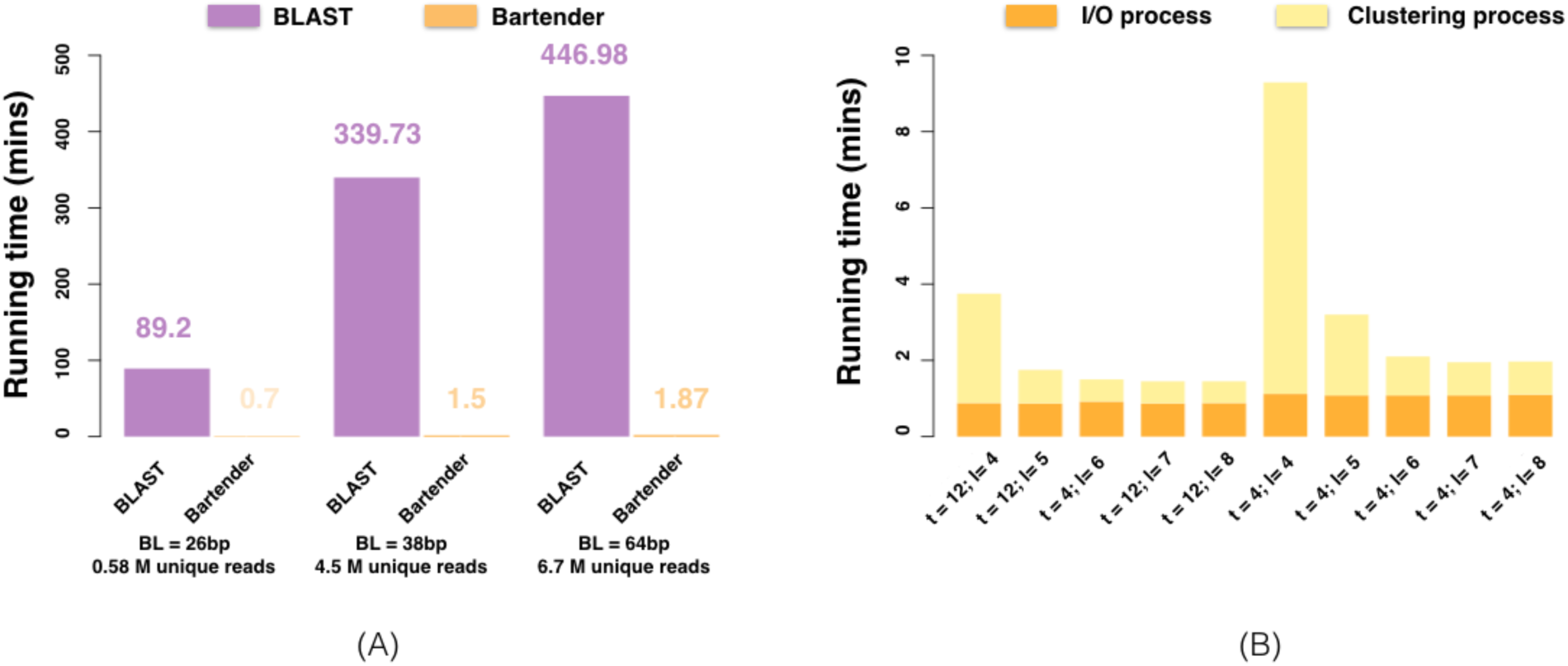
Bartender speed. (A) Running time for Bartender and BLAST on simulated data of barcodes of different barcode lengths (BLs), using 12 threads (t = 12). Bartender was performed with a seed length of 5 (l = 5). B) Bartender performance of simulated 38 bp barcodes using a variable number of threads (t) and seed lengths (l). Input/output (I/O) time is shown in orange and clustering time in shown in yellow. For t = 12, a desktop equipped with 3.5 GHz 6-core intel xeon E5, 64 GB memory was used. For t =4, a laptop equipped with 3.0 GHz Intel core i7 and 16GB memory was used.

### Bartender prevents over-clustering

To make merging decisions between reads/clusters, Bartender uses a modified two-sample proportion test that considers both cluster distances and cluster sizes (see Methods). To determine the cluster distances, Bartender uses Hamming distance (see Methods) instead of Levenshtein distance for two reasons. First, the time complexity of computing Hamming distance is linear, while that of Levenshtein distance is quadratic, in barcode length, so Hamming distance is more efficient for long barcodes. Second, insertions or deletions (indels) are generally rare on Illumina sequencing platforms (our observations), so the accuracy gained by using Levenshtein distance is limited. Theoretically, indels can make Bartender underestimate cluster sizes, however we did not observe this with real data (see below). Regardless of the method used to calculate cluster distances, a major caveat of making merging decisions based solely on sequence similarity is that distinct barcodes that are close in sequence will be erroneously merged. To overcome this problem, we take advantage of the fact that for two clusters that are close in distance, if one cluster contains true barcode reads and the other contains sequencing errors of this barcode (i.e. the two clusters arising from the same barcode), then the cluster containing errors is expected to be at a much lower frequency that is a function of the error rate and the barcode length (see Methods). If, however, each cluster represents a true barcode, then these two clusters are more likely to be at similar frequencies. Therefore, Bartender considers the relative frequencies of two clusters in making merging decisions, with clusters that have larger frequency disparities being more likely to be merged.

To determine the accuracy of Bartender, we generated a simulated data set that consists of 100,000 random 20mer barcodes, with their frequencies following an exponential frequency distribution (mean = 100) that yields ~10M total reads (Supplementary Table S1, Figure S1). The simulation included a 2% combined per nucleotide error rate of PCR and sequencing and yielded ~4.8M unique reads. Using both Bartender and BLAST, we output the consensus sequence (majority vote at each nucleotide position) and the count of each predicted barcode (cluster). Predicted cluster counts were compared to the true counts known from the simulation. For this comparison, clusters with less than three reads were ignored because these low-frequency clusters are more likely to be erroneous and derived from another cluster in the pool (see Methods). Using the remaining 96,610 clusters, the two methods found nearly the same number of clusters (96,517 for Bartender, 95,537for BLAST), however Bartender generally estimated the true count with greater accuracy (Figure 2). In particular, many BLAST cluster estimates were too large, indicating that closely-spaced clusters were erroneously merged (Figure 2, green triangles). BLAST over-merging resulted in a higher number of false negatives (Bartender = 93, BLAST = 1073), and marginally less false positives (Bartender = 94, BLAST = 53), with Bartender errors exclusively occurring in small clusters. We note that over-merging is not a problem that is unique to BLAST. All previous methods that use only a distance-based similarity threshold (e.g. 2 mismatches) to merge similar reads, but not cluster size information (unique to Bartender), are subject to the same over-merging artifacts as BLAST.

**Figure 2.**
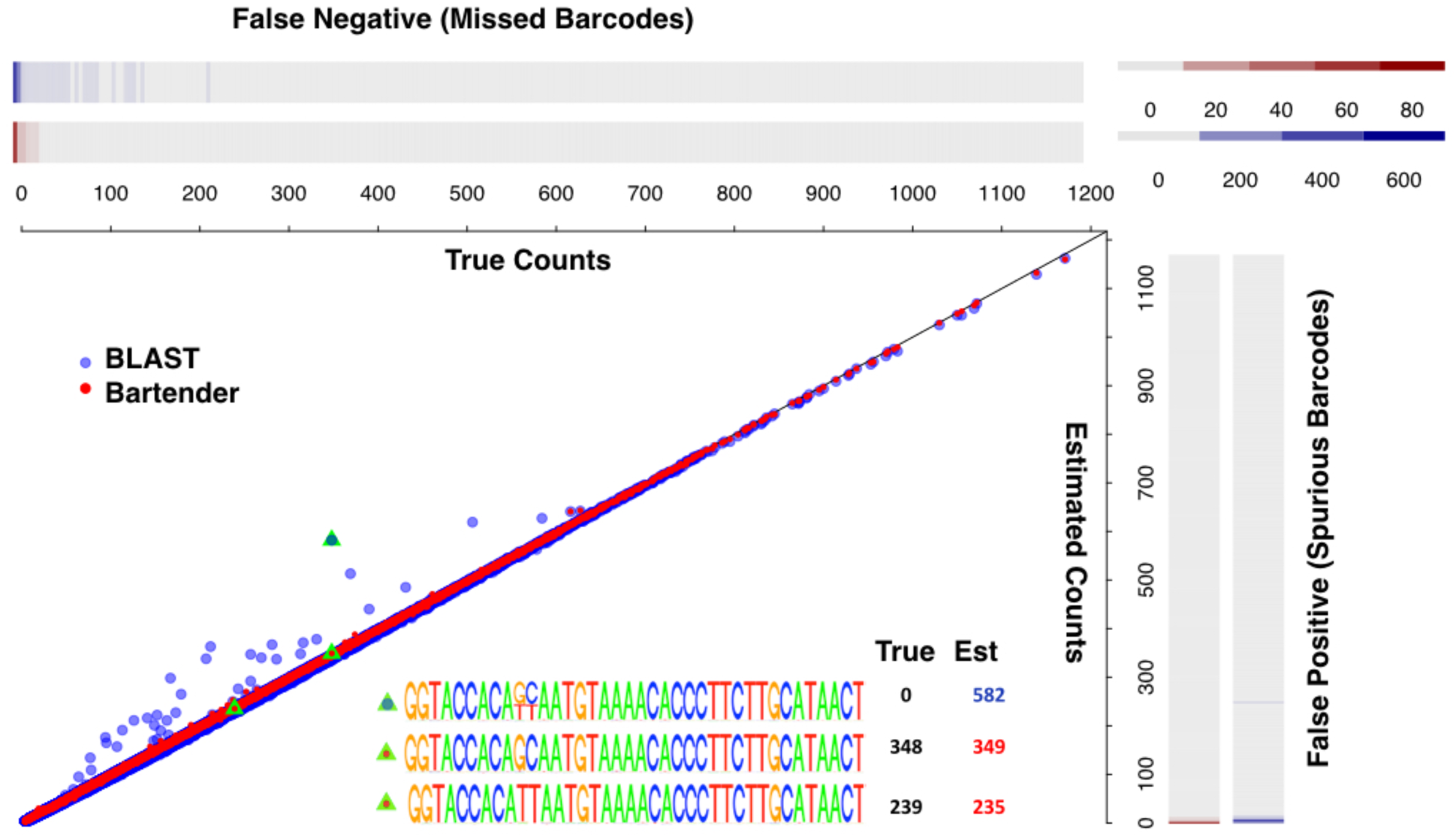
Comparison of Bartender and BLAST on simulated data. A scatter plot of the estimated and true counts of each barcode cluster for Bartender (red) and BLAST (blue). Over-clustering by BLAST but not Bartender results in some unique barcodes being merged. Green triangles and inset of position weight matrices show a representative example of BLAST over-merging of barcodes that are two mismatches away from each other. Heatmaps show the rate of false positives and false negatives for Bartender (red) and BLAST (blue).

### Bartender errors and sequencing depth

Barcodes with low sequencing depths are difficult to accurately count for several reasons. First, sampling noise (derived from which molecules happen to be sequenced) scales with the square root of the expected number of reads (see Methods). That is, barcodes with less reads will have high coefficients of variation simply because of sampling on the sequencer. Second, small read numbers can often result in an erroneous cluster center (the predicted barcode sequence does not match the true barcode sequence) because PCR or sequencing errors in one read would have a large impact on the predicted center. These errors in small clusters often result in both false positives and false negatives because reads matching a true barcode sequence are missing while those matching a new (erroneous) barcode sequence are present. Third, merging decisions between small clusters become less accurate. As described above, Bartender takes advantage of cluster size information to make merging decisions. An unequal frequency between clusters favors merging because it becomes more likely that the low frequency cluster represents a sequence error of the high frequency cluster. However, when a barcode is present at low frequencies, both true reads and errors are expected to have few counts, so differences in frequency become blunted and less informative.

To examine how barcode sequencing depth impacts Bartender accuracy, we simulated two data sets containing 100,000 true barcodes and ~10M reads, with error rates of 2% (Figure 3A) and 0.33% (Figure 3B), respectively. In both cases, Bartender clustering introduced almost no errors for moderately large clusters (>10 reads). Counting errors for these large clusters are due almost exclusively to sampling at the sequencer. For smaller clusters (<10 reads), the clustering process did introduce additional error for reasons described above. However, the additional error due to clustering is small, especially at the lower 0.33% sequencing error rate, which is also the rate observed in real data (supplementary figure S2). Nevertheless, sequencing at a coverage sufficient to read each barcode at least 10 times would virtually eliminate Bartender errors.

**Figure 3.**
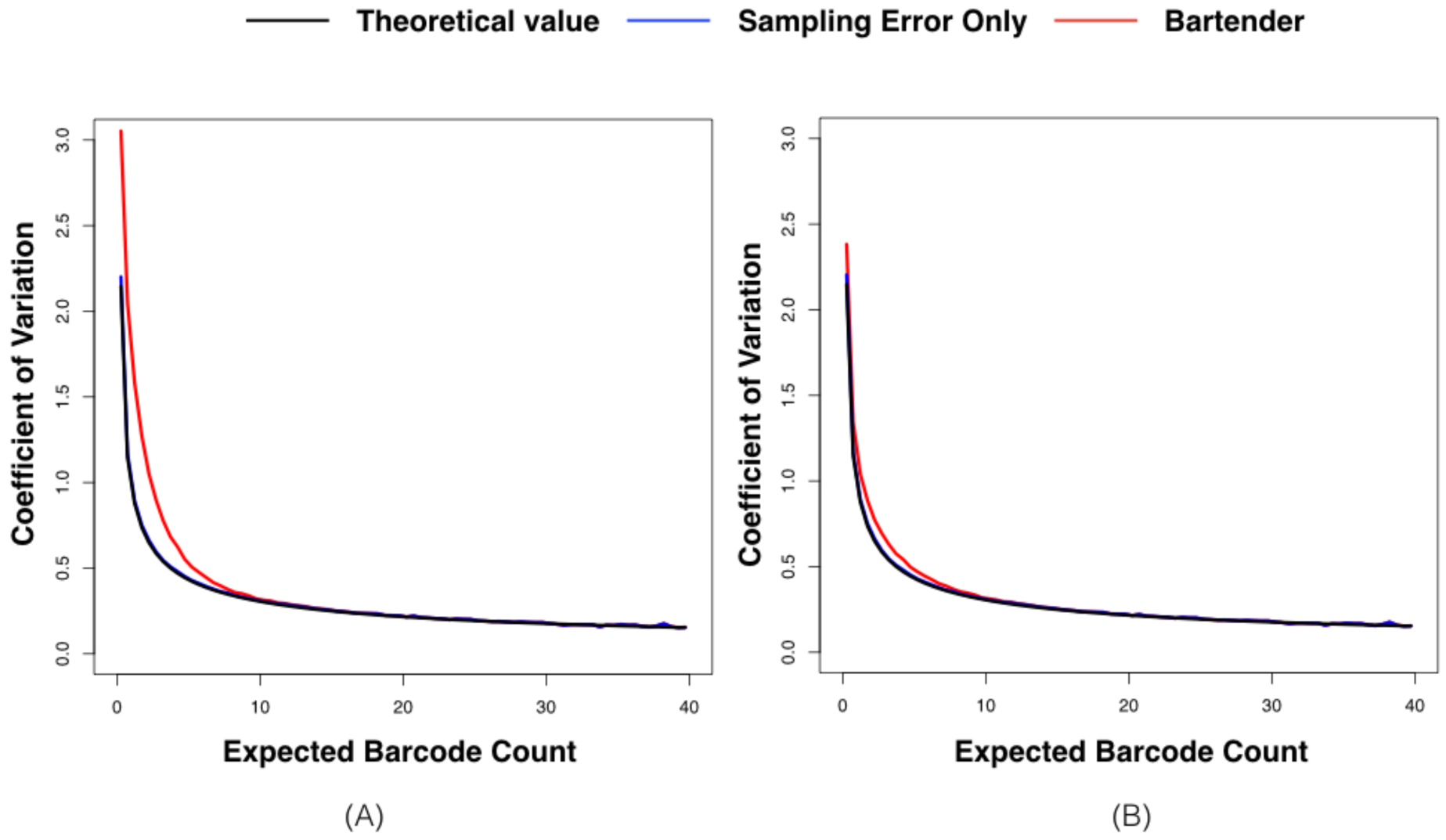
The impact of sequencing depth on Bartender performance. A plot of the barcode count by the coefficient of variation (CV) for that count on simulated data with a 2% (A) and 0.33% (B) combined error rate of PCR and sequencing (sequencing error). The black lines are theoretical values, which follow the Binomial distribution. The blue lines are the C V of sampling (at the sequencer) alone, without sequencing errors or errors introduced by Bartender clustering. The red lines are the CV after running Bartender, and include sampling, sequencing, and clustering errors. All lines are smoothed with window size 0.5.

### Application to high-complexity bar-seq data

To compare Bartender and BLAST performance on real sequencing data, we next clustered published data of a high-complexity barcode library (~0.5M barcodes, 20 random bases, 26 total bases) that has been sequenced deeply (~136M reads) to generate ~3M unique barcode reads (Levy et al., 2015). The two methods discovered 488,983 overlapping barcode clusters, with 2,602 additional clusters identified only by Bartender and 22 only by BLAST (Figure 4A). Of the 2,602 Bartender-only clusters, 2,301 contained more than 10 reads (Figure 4B), indicating that they are likely to be true barcodes (from the Bartender error analysis above). The 22 BLAST-only clusters were all present at low frequencies (<5, Figure 4B), suggesting they likely arose from read errors. The overlapped clusters between BLAST and Bartender (Figure 4D) reveals that the two methods generally correlate well (Figures 4D and S3, Pearson’ correlation = 0.97). For many barcodes, however, BLAST estimated higher counts than Bartender, echoing findings on simulated data (Figure 2), and suggesting BLAST is over-clustering. To investigate this possibility, we examined these outliers and found that, in most cases, BLAST merged two or more distinct barcodes together. A representative example of BLAST over-merging is shown in Figures 4C and 4D (grey triangles): several barcodes with similar counts that are close in sequence space can be distinguished by Bartender but not BLAST. Similar speed improvements found with simulated data are also found with this real data set (Bartender = 5.4 minutes, BLAST = 337 minutes, 4 threads). A detailed examination of Bartender running time with this real data is given in Figure S4.

**Figure 4.**
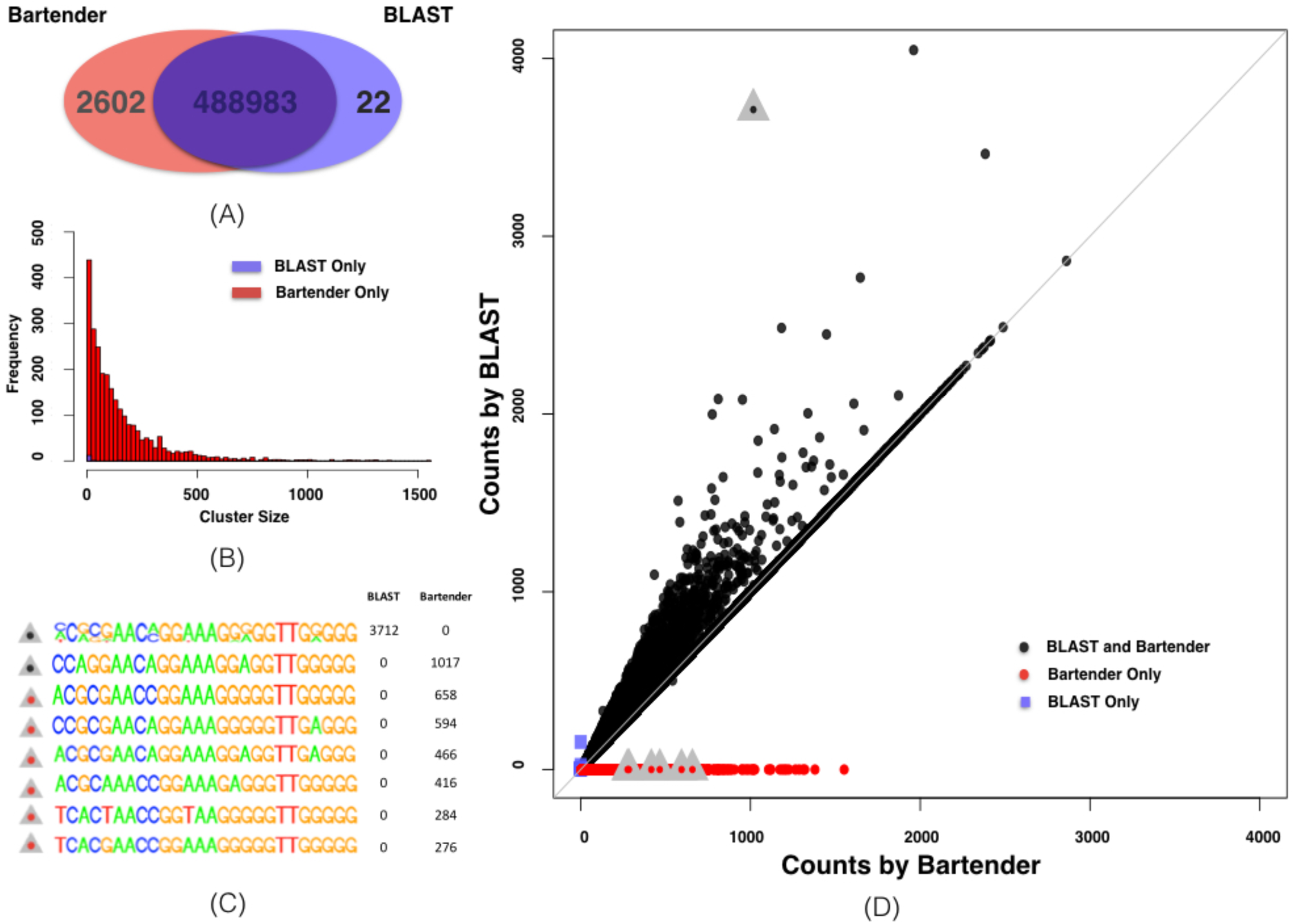
Bartender and BLAST performance on real data. (A) Ven diagram of the number of clusters identified by Bartender and BLAST. (B) Histogram of the number of counts for barcodes identified by BLAST but not Bartender (blue), and Bartender but not BLAST (red). Some large Bartender clusters (size > 1500) are not shown. (C) Position weight matrices of the highlighted clusters from (D). The first cluster is generated by BLAST, is variable at multiple nucleotide positions, matches the sequence of the first Bartender cluster (second black dot), and incorporates many additional unique clusters that are distinguished by Bartender (red dots). All Bartender clusters display low variation at each nucleotide position. D) Scatter plot of the counts of each barcode detected by both Bartender and BLAST (black), Bartender only (red), and BLAST only (blue squares). Grey triangles are a representative example of BLAST over-clustering and are further explained in (C).

### Bartender includes UMI handling

Unique molecular identifiers (UMIs) are additional, usually random, sequences that are added to template molecules before PCR that allow an investigator to detect and remove PCR duplicates and thereby improve the accuracy of amplicon counting (Kivioja et al., 2012; Levy et al., 2015). Bartender allows a user to attach a UMI sequence to each barcode prior to clustering: the user must simply generate a comma-separated file that contains one barcode and one UMI on each line. Following clustering, Bartender will search for identical UMIs within each cluster and report counts that include or exclude repeated UMIs (putative PCR duplicates). We note here that UMI length must be carefully considered as part of the experimental design, and provide more some general guidelines in the discussion.

### Bartender includes multiple time point handling

A growing number of studies perform time course bar-seq experiments that require tracking barcode lineages over time (Gresham et al., 2011; Levy et al., 2015). In some cases, lineages will persist at low frequencies or be driven over time to extinction. These sorts of scenarios present additional challenges for Bartender clustering. As discussed above, a small cluster is more likely to result in a difference between the true barcode sequence and the sequence Bartender calls. These cluster center errors will result in some barcodes at low frequencies artificially “disappearing” at some time points and then “reappearing” at others, greatly impacting interpretation of the lineage trajectory. One work-around we previously employed (Levy et al., 2015) was to pool reads from all time points, cluster the big pool to build a map between each unique read and a barcode cluster, and then reconstruct the counts at each time point from this map. However, this approach is computationally expensive and can easily exceed the memory limitations of even a powerful desktop computer. To avoid these issues, we include a “multiple time point” mode in Bartender. This feature allows time points to be clustered independently, thereby reducing computational overhead and allowing for parallelization. Once individual time points are clustered, Bartender joins clusters between adjacent time points under the assumption that barcodes in a time point are a subset of the clusters the previous time point. For a time point *t*, Bartender first joins clusters with the exact same center (sequence) to *t*−1. Then, any unjoined clusters in *t* are joined to clusters in *t*−1 if their centers that are within one mismatch. This second step (joining across mismatches) is particularly important for barcodes that are being driven to low frequencies. For these small clusters (e.g. <3, see above), the estimated cluster center is more likely to be erroneous, and by using information from the prior time point, Bartender becomes more accurate at estimating their frequencies.

To test the accuracy of the multiple time point feature, we simulated an evolution of 100,000 barcoded cells over 112 generations (Supplementary Materials). Most of these cells were set to have an identical fitness which we define as 0. However, 5000 (5%) cells were set to have a higher fitness that is drawn from a truncated exponential distribution *(λ =* 0.00083 and bounds of 0.05 of 0.15). We assume the fitness of each cell lineage is unchanging over the course of the competition (no adaptive mutations). Over time, higher fitness lineages will expand and drive lower fitness lineages to low frequencies and eventually to extinction. We simulated sampling barcode lineage frequencies at various time points, assuming ~10M total reads per time point, performed Bartender clustering, and compared these results to the true barcode frequencies known from the simulation (Figure 5A). Additional time points generally improved Bartender’s performance. Bartender detected 100,389 barcode clusters including all true barcodes (no false negatives) and 389 erroneous barcodes (false positives). All false positives were only present in the first time point and no subsequent time points, and would be easily identified and ignored by an investigator. Bartender was extremely accurate at estimating barcode frequencies over time, but accuracy suffered when lineages fell to low frequencies of 2-3 reads per barcode. To exemplify this point, we sampled the barcode frequencies more densely during the mass extinction event that begins at ~70 generations in our simulation (Figure 5B). During this time, we find that Bartender slightly underestimates the number of barcode lineages that are still present in the pool. As discussed above, these errors are generally due to PCR and sequencing errors in low frequency lineages that cause Bartender to miscall barcode cluster centers. When the number of lineages becomes stable (~5000 large lineages at 81 generations), Bartender is again able to correctly identify all barcodes that exist in the pool.

**Figure 5.**
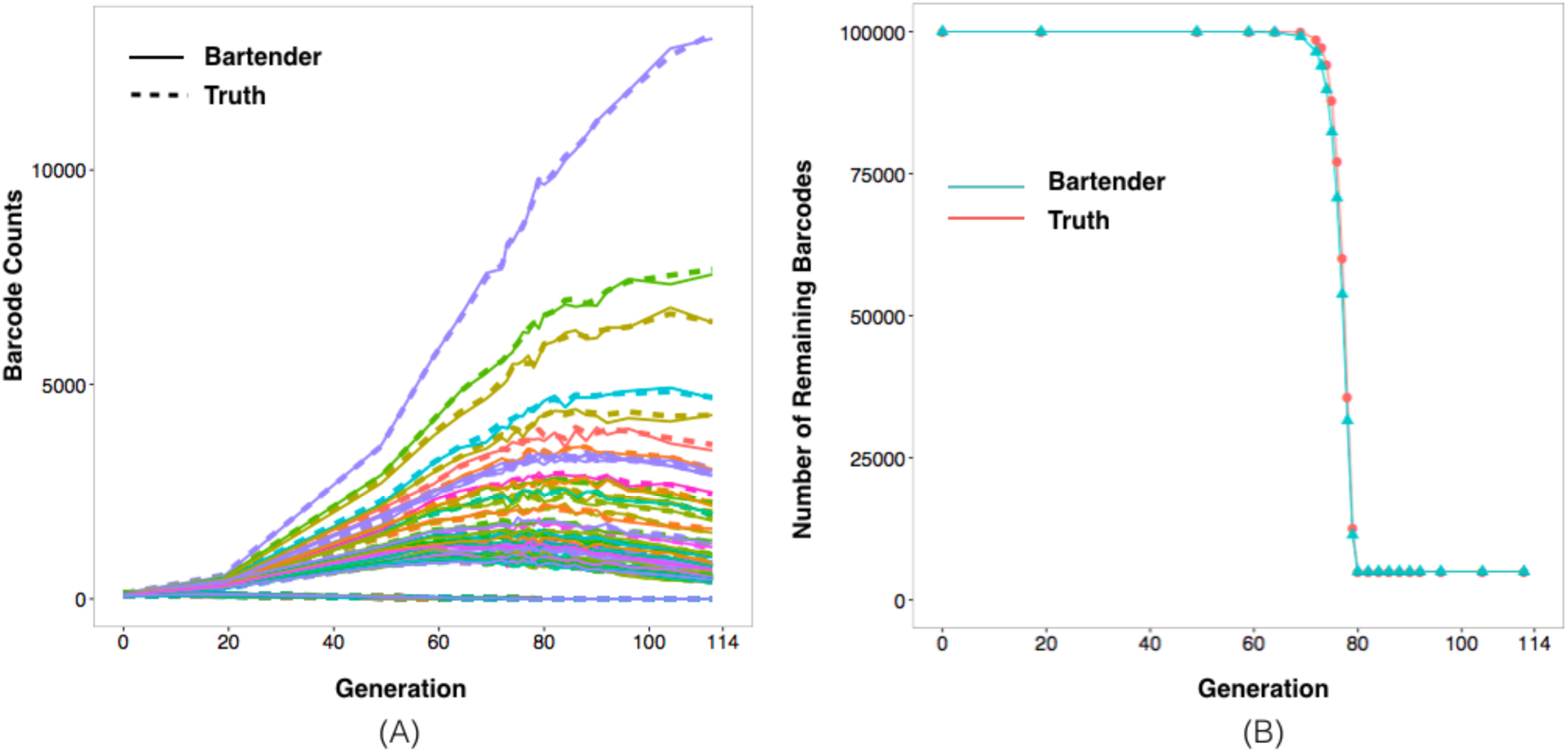
Bartender performance on time course data. A simulation was performed of 100,000 barcoded cells with different fitness coefficients that are evolved in competition for 112 generations (A) Lineage trajectories of 1000 randomly selected barcodes (colors). Solid lines are trajectories estimated by Bartender and dashed lines are the true trajectories. (B) The number of barcodes present in the pool at a count greater than 1 (red) and the number detected by Bartender (turquoise). At ~70 generations, lineages without a growth advantage begin to go extinct. During the following mass extinction, Bartender slightly underestimates the number of barcodes present.

## Discussion

To the best of our knowledge, Bartender is the first general clustering algorithm for accurate and fast counting of barcode and amplicon reads. Bartender’s accuracy stems from a new statistical test schema that uses both nucleotide sequence and cluster size information to prevent overmerging; its speed stems from a novel binning strategy and a computationally efficient greedy clustering algorithm. Bartender includes handling of both UMIs and time course data, and promises to be a useful tool for a large number of diverse applications.

Bartender includes three parameters that are tunable for different applications: the maximum cluster distance that may be merged (*d*), the seed length (*l*, see Results), and the merging threshold (*z*, see Methods). We recommend using the default seed length (*l* = 5) and maximum cluster distance (*d* = 2) for applications where shorter run times are not a priority. The remaining parameter, *z*, should be set according to the expected coverage per barcode and the barcode library complexity. For low to medium coverage (<500 reads/barcode), we recommend starting with the default setting for *z* (= 5). However, in some cases, it may be necessary to adjust the *z*. For example, we have noticed some nucleotide sequences in our random barcode libraries are more prone to PCR or sequencing errors (i.e. errors at that position are greater than *e*), and, if these errors occur within an abundant barcode, they may cause the same erroneous read to occur at a high frequency and therefore be interpreted as an independent barcode cluster. At extremely high coverage (e.g. >10,000 reads/barcode in a low-complexity barcode library), this problem is amplified because a high error position in any barcode will create this artifact. In this case, we recommend setting *z* higher. In cases where all barcodes are expected to be distant (an average of 5-6 mismatches from a nearest neighbor), we recommend disabling the merging threshold (*z* = −1) to make merging decisions based on the cluster distance (*d*) only. In this scheme, all sequences within a user specified distance of each other will be merged, however, because barcodes are distant, it is unlikely that any real barcodes will be merged together (over-merging). Furthermore, disabling the merging threshold will remove the possibility of nucleotide sequences with abnormally high errors causing artifacts. We note that Bartender will only perform well if most barcodes within the pool are sufficiently spaced such that they are at least 3-4 mismatches away from a nearest neighbor (see (Blundell & Levy, 2014; Levy et al., 2015) for a discussion on random barcode design). In cases where this rule is broken (Bhang et al., 2015), Bartender performance will be encumbered, as will any existing clustering approach.

Bartender speed is mainly due to the fact that it partitions unique reads into bins and restrains sequence comparisons to those within each bin. We use the entropy at each nucleotide position to prioritize seeds that have the highest probability of creating a maximal number of bins, and thereby minimizing the number of pairwise sequence comparisons. Barcode designs with regions of varying complexity (e.g. (Goodman et al., 2013; Kosuri et al., 2013; McKenna et al., 2016)) could potentially disrupt this process and greatly slow Bartender. Take, for example, a barcode that is split into low complexity (e.g. 1000 random variants) and high complexity (e.g. 1M random variants) regions. Because a nucleotide in either region would be expected to have high entropy, Bartender may choose initial seeds that contain only low complexity nucleotides resulting in far fewer bins. One potential solution would be to use a seed selection protocol that considers associations between nucleotides (e.g. information gain or mutual information), however this is not implemented here.

Bartender removes PCR duplicates by simply searching for repeated UMIs within each cluster (exact matches) and removing these from the counts. Because it searches only for exact UMI matches, a PCR or sequencing error that happened to occur in a repeated UMI would not be recognized as a repeat and thus result in over-counting of that cluster. However, the alternative, merging UMI with similar sequences, raises greater problems. Large clusters may contain many UMIs, and, because UMIs are generally short, UMI clustering would erroneously merge many distinct UMIs that are close in sequence. Even using our exact match criterion, it is possible that extremely large barcode clusters will begin to use up all available UMIs resulting in undercounting. For example, a barcode that is read 100,000 times and contains an 8mer UMI (4^8^ = ~65,000 possible sequences) will necessarily have UMI repeats even when each sequenced read stems from a unique template molecule. To avoid these problems, we recommend selecting a UMI length that results in at least 10-fold more possible sequences than the largest expected cluster.

Bartender includes a multiple time point mode that is designed to generate barcode trajectories for time course data. Importantly, information from a previous time improves count estimates for low frequency barcodes because erroneous cluster centers (due to sequence errors) are more likely to be assigned to a true barcode. This mode assumes that all barcodes at any time point are a subset of the barcodes in the previous time point. In some cases, this mode could be applied to experiments other than a time course (e.g. bar-seq across different conditions). However, we advise extreme caution. If there are only a few (2-3) conditions, and there is a large overlap in the barcodes present in each condition, then multiple time point mode could be directly applied using an arbitrary order of the conditions. However, in cases where the overlap between barcodes present in each condition is expected to be small, we do not advise using the multiple time point mode.Rather, Bartender should be applied to each condition separately and then barcode clusters from different conditions should be compared by another method.

Another useful feature of Bartender is that it reports an entropy-based measurement for cluster quality to an output file, which can be used in downstream analyses as an indicator that a cluster might contain more than one true barcode. The cluster quality is a measure of the read heterogeneity within a cluster. For each cluster, a position weight matrix is generated from all the unique reads contained in the cluster and the cluster quality is reported as the largest binomial entropy value across all positions in this matrix (see Methods).

Based on our experience, Bartender will work well with a standard laptop or desktop computer for most applications. Available memory (RAM) is generally the limiting factor for Bartender processing, and the necessary RAM is a function of the number of unique barcode reads and the barcode length. We recommend 4-6 GB RAM for datasets with less than 1M unique 40mer barcodes, and 8-10 GB RAM for datasets with less than 3M unique 60mer barcodes.

## Methods

### Dissimilarity measures between reads and clusters

Suppose there are *N* unique reads and let *r_i_* be the *i*th unique read with a frequency of *f_i_*. For two distinct reads (*r_i_* and *r_j_*) with the same length *L,* their *Hamming distance* (Hamming, 1950) is defined as follows:

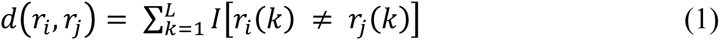

where *r_i_*(*k*) is the nucleotide at the *k*th position of sequence *r*_i_, and *I*(*x*) is an indicator function such that

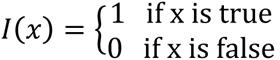

For a particular barcode cluster *C* that may contain several unique reads, its size is defined to be:

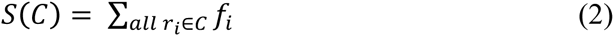

The centroid of the cluster *C* is defined as a sequence *c* = [*c*_1_*c*_2_ … *c_L_*], where 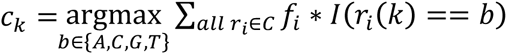 for *k* = 1, …, *L*. That is, the centroid consists of nucleotides that are most frequent at each position. Since a cluster usually corresponds to a barcode, the centroid can also be viewed as an estimate of the corresponding barcode.

For two distinct clusters *C_x_, C_y_*, let *c_x_, c_y_* denote their centroids. We use *Hamming distance* between the two centroids as the dissimilarity metric to define the distance between the two clusters, which is given as follows:

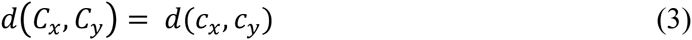

### The BLAST clustering method

In the BLAST-based clustering protocol, three major steps are critical for its accuracy and efficiency. The first step is to find a size threshold to split unique reads into high-and low-frequency groups, where high-frequency reads are likely to be a true barcode and low-frequency are likely to be sequencing and/or PCR errors. This splitting step is necessary to reduce the number of pairwise sequence comparisons and improve computational speed (clustering can take >10-fold longer without the splitting). The size threshold is related to the underlying experimental design (i.e. the expected number of barcodes and their underlying frequency distribution) and the sequencing parameters such as sequencing depth and error rate. In the simulated and real (Levy et al., 2015) data sets we use, the size thresholds are set to 2 and 10, respectively. Once the unique reads are divided into two lists, the second step is to group similar high frequency sequences by performing all pairwise comparisons within the high-frequency list, where the similarity is a function of the e-value in the BLAST package. Two clusters are joined if any two sequences (one in each cluster) pass the e-value threshold. Cluster joining continues until the number of clusters is stable. Lastly, sequences with low-frequency unique reads (below the pre-specified size threshold) are then matched to the high-frequency clusters by BLAST using the same criteria. A majority vote method is used at each nucleotide position to define the cluster center, which represents a cluster tag for any downstream analysis.

### Bartender workflow overview

Bartender processes barcodes with different lengths independently. The overall Bartender workflow for processing barcodes with same length is shown in Figure 6. First, total reads are tabulated to form a list of unique reads and corresponding frequencies. Using this list of unique reads, a sequence of overlapping 5-8 base pair seeds is selected. These groups of unique reads are considered as initial clusters, and are partitioned into bins using the first seed. Clusters within each bin are merged using a statistical test described below. These new clusters then serve as the input for the next round of binning, and this is repeated until all seeds have been processed. Lastly, three output data files are generated: the consensus cluster sequence (the barcode) and its counts, the quality of each cluster, and a map between each unique read and the cluster it belongs to. Unique Molecular Identifiers (UMIs) for each read that are stored with each cluster are then used to remove PCR duplicates by searching for identical UMIs within each cluster. The detailed logic is described by Algorithm 1 in the supplement.

**Figure 6.**
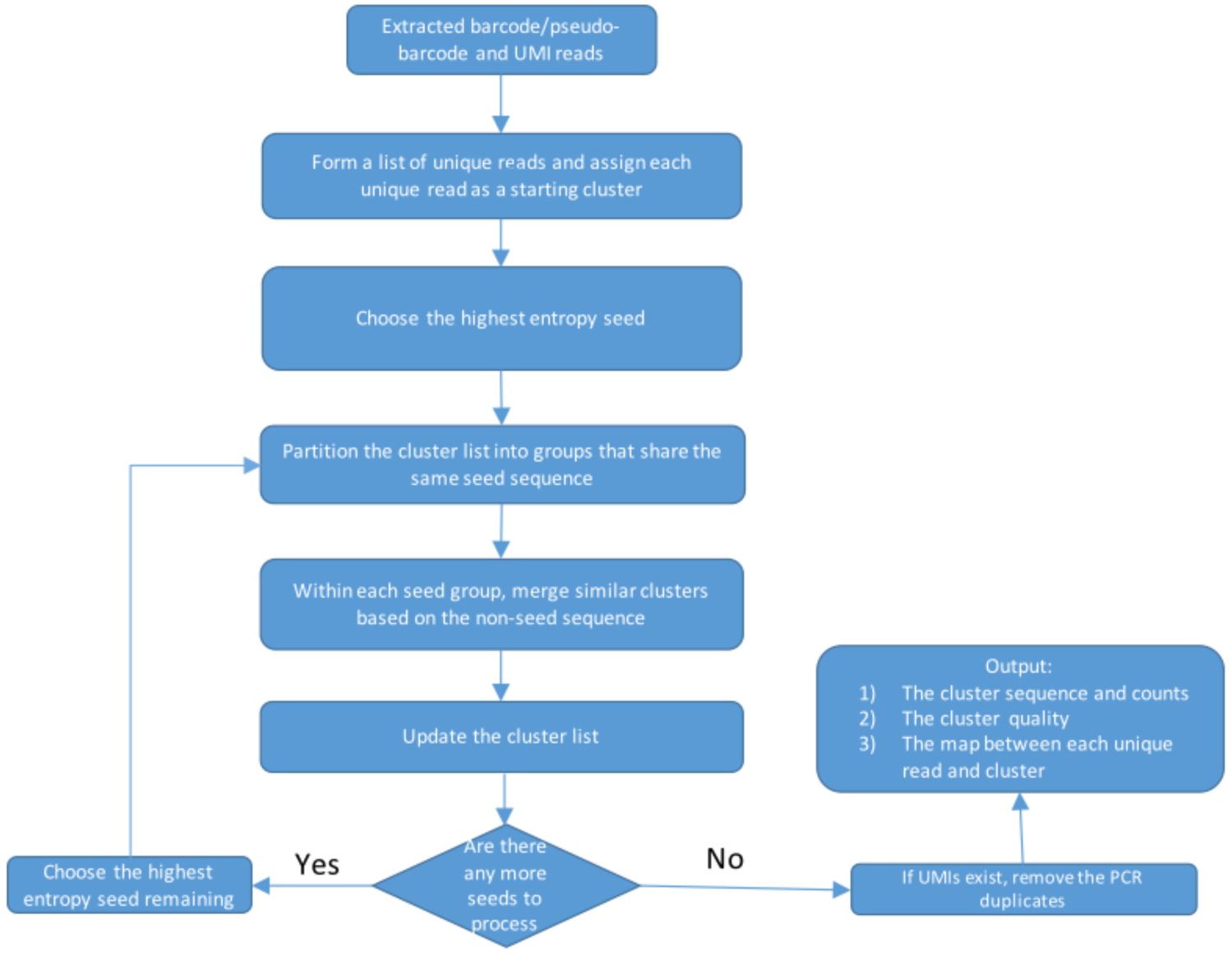
A schematic flowchart of the Bartender clustering algorithm.

### Bartender Extractor

Bartender extractor is a simple command-line tool designed to extract barcode from raw reads. It assumes that each read has at most one barcode that is flanked by some fixed sequences to locate the barcode. A barcode template could be one large random region or consist of multiple small random regions separated by spacers. Because indels may occur during barcode generation, users can specify a length range for each random region. While extracting, this tool keeps track of the total number of base errors identified in the flanking (constant) regions and estimates the sequence error using the percentage of these errors. The extractor allows at most one mismatch in each flanking region, so, in order to obtain relatively accurate error estimation, the length of a flanking sequence should not be set to be too long (we recommend 5 bases). If sequence error is not uniform across all positions in an amplicon and error biases in the flanking regions exist, this sequence error estimate might be slightly incorrect. However, we recommend using this error rate as the baseline for Bartender clustering. The Bartender clustering command provides a second estimate of the error rate based on the random regions. If the error rate estimated by the clustering component is significantly different from that estimated by the extractor, we recommend rerunning Bartender clustering with more conservative parameter values (i.e. with smaller *d* and/or *z*).

The extractor transfers the specified barcode template into regular expression and extracts the matched subsequence from each read in the raw file (FASTQ or FASTA format). A user defined quality threshold can also be used to filter out low-quality barcodes (only applicable to FASTQ format). When the average quality is below this threshold, these reads will be ignored. The matched sequence (flanking regions removed) and the corresponding line number in raw (FASTQ or FATSA) file are output to a file. This file can be used directly for Bartender clustering. Alternatively, this file can be used in combination with user-generated UMI file, to replace the line number with the UMI prior to clustering. If UMIs are included, Bartender will by default remove duplicates during clustering.

### Seeds selection and binning

To calculate entropy values for N unique reads with length L, we first summarize a position weight matrix, i.e. the frequency of each of the four nucleotides (A, C, T, G) at each position: 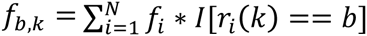, where *b* ∈ [*A, C, G, T*] and *k* = 1, …,*L*. At each position *k*, the four nucleotides are dichotomized into two groups, one with the most frequent nucleotide (the major allele) and another with the other three nucleotides (the other alleles). The relative frequencies of the major and other alleles are then calculated and denoted as 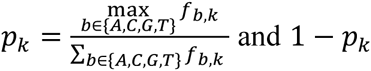 respectively. The entropy at position k is defined as:

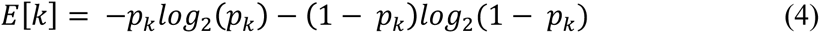

Seeds are then selected based on the order of the entropy values. Positions with larger entropies are used in earlier iterations of binning. Nucleotide positions with too low entropies below a prespecified (or user-defined) threshold are excluded from seed selection.

Next, clusters, based on their centroids, are sorted into different bins based on selected seeds. The binning process is demonstrated in Figure 7. The centroids of all clusters in one bin contain the same seed sequence. There are two major computational advantages of binning by seeds. First, unnecessary pairwise comparisons between distant clusters are dramatically reduced because only clusters within the same bin will be compared. Second, each bin can be processed independently, making it easy to parallelize the algorithm on multiple-core computers. Intuitively, a longer seed generates more bins and correspondingly decreases the number of clusters in each bin. The seed length is a critical parameter that balances the speed and accuracy of the clustering algorithm. That is, longer seeds increase speed by reducing pairwise comparisons, but are more likely to leave spurious barcodes ungrouped, thereby increasing false positives. By default, seed length is set to be five (see Figure 7 for details), and Bartender iterates through the seed position list using a sliding window with a size of the seed length and slide one nucleotide at a time.

**Figure 7.**
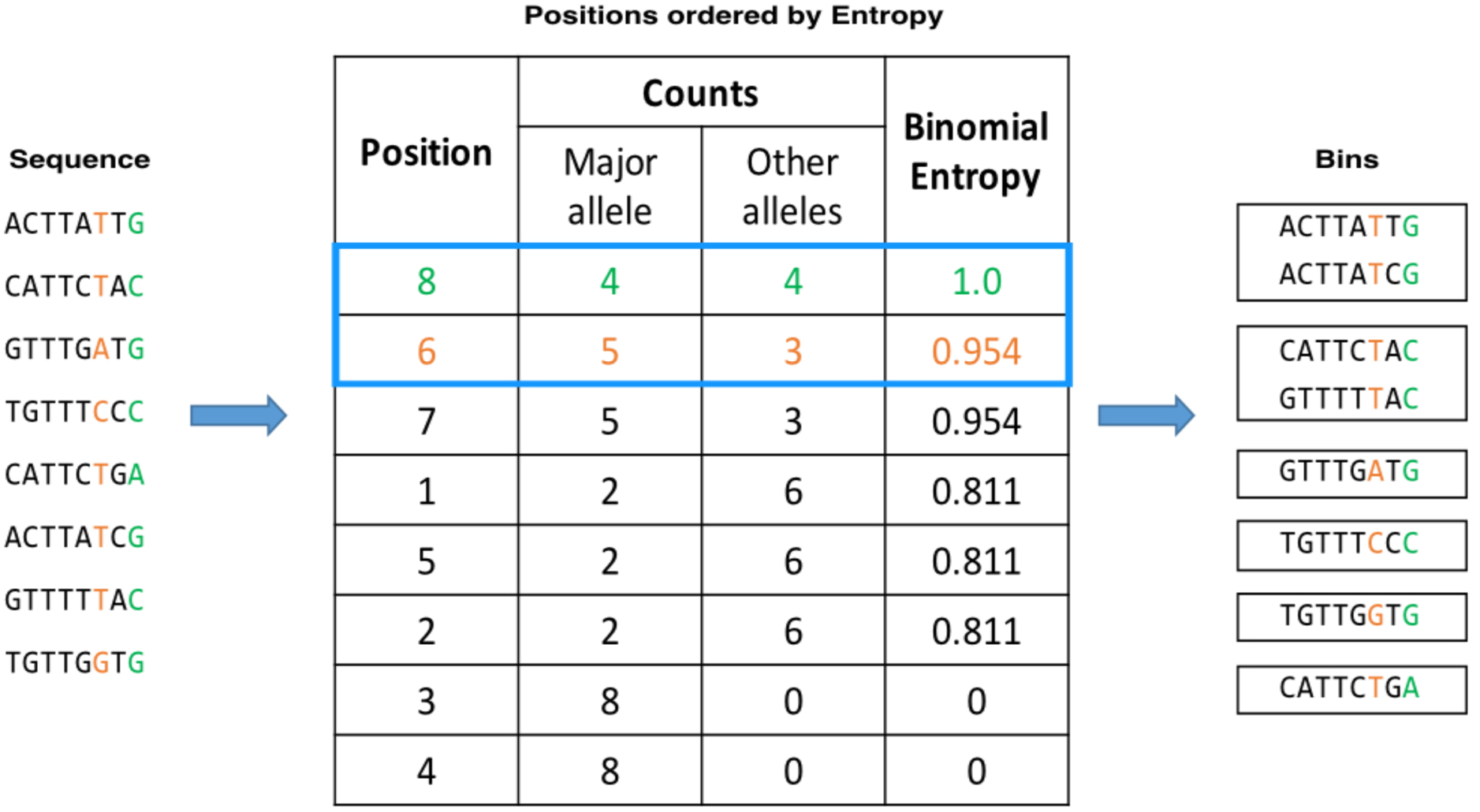
Binning by seeds. A toy example of seed selection on eight unique sequences with a seed size of two. The entropy value at each position is calculated and positions are sorted in descending order, with ties broken arbitrarily. Here, positions eight and six have the highest entropy and are used as the first seed, breaking the sequences into six bins (right). The next seed is positions are by default positions six and seven.

### Clustering within one bin

Since there may be thousands of barcode clusters within one bin, pairwise comparisons within a bin are still computationally expensive. To further improve the speed, we developed a greedy clustering algorithm that uses cluster frequency information to prioritize comparisons. The rationale is that clusters with high frequencies are more likely to represent a true barcode, while those at low frequencies are more likely to be errors. We take advantage of this and adopt a strategy similar to that used in the BLAST method by first splitting clusters into high- and low-frequency bins based on the mean cluster size, which is derived from the empirical distribution of the sizes of all clusters. Since error reads are expected to greatly outnumber reads with true barcode sequences, the mean cluster size is expected to partition the majority of error-containing sequences (and maybe a minority of true barcodes) into the low-frequency group. Each cluster in the low-frequency group is compared to clusters in the high-frequency group using the statistical test described below, and three scenarios may occur: if no high-frequency cluster is found, this low-frequency cluster is added to the high-frequency cluster group, as it is likely to be a true barcode cluster; if only one high-frequency cluster is found, the low-frequency cluster is merged to this cluster; if >1 high-frequency clusters match, the low-frequency cluster will be merged with the high-frequency cluster with the closest sequence. If multiple high-frequency clusters are equally close to this low-frequency cluster, then it will be merged into the largest high-frequency cluster. Following this step, pairwise comparisons between all high-frequency clusters are performed using the same statistical test described below, and clusters are merged if they pass. A more detailed description is given in the supplement Algorithm 2.

### Cluster merging

Clusters that are close in distance may arise from two possibilities: (1) one cluster is a true barcode cluster and another is a cluster with error reads, and (2) both clusters are true barcode clusters. To avoid over-merging in the second case, we use a modified two-sample proportion test to make the merging decision for the two clusters.

Let *C*_1_ and *C*_2_ denote the two clusters being tested, with centroids of *c_1_* and *c*_2_, respectively. The distance between *C*_1_ and *C_2_* is *d*(*C*_1_,*C*_2_) = *d*(*c*_1_,*c*_2_) = *d*. Let *C*_3_ denote the new cluster when *C*_1_ and *C*_2_ are merged, with a new centroid *c*_3_, which is the center of the larger cluster between *C*_1_ and *C*_2_. The sizes of *C*_1_ and *C*_2_ are S(*C*_1_) and *S*(*C*_2_), and the size of *C*_3_ is *S*(*C*_3_) = S(*C*_1_) + *S*(*C*_2_). Let *e*_i,3_,*i* = 1,2 be the cumulative number of base pair errors of reads in *C_i_* with respect to the new cluster *C*_3_:

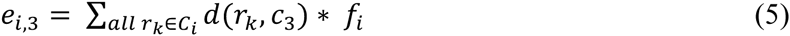

Let *p_i_*_, 3_ be the error rate of cluster *C_i_* with respect to the new cluster *C*_3_, which is defined as:

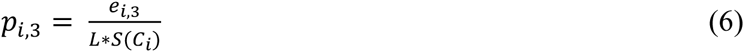

The idea is that if one barcode cluster indeed originates from errors of another barcode cluster, the size of the error cluster should be much smaller than that of the true barcode cluster.

The hypotheses can be formulated as

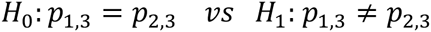

The test statistic is given by:

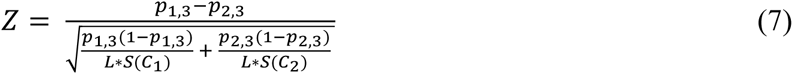

Z becomes large under two scenarios: 1) the distance between cluster centroids is large, or 2) there is a small but consistent sequence difference between two sufficiently large cluster centroids. Therefore, we reject *H*_0_ for large Z values when two clusters should not be merged.

### Evaluation of the Bartender accuracy

Differences between the true barcode size (count) and the Bartender estimate result from three primary sources of error: sampling error, sequence error, and additional error introduced by Bartender clustering. Sampling error refers to the size difference between the theoretically expected copy number (based on the relative frequency in the cell population) and the number of copies that are actually sequenced on the sequencer. This error cannot be removed and thus represents the lower bound of all error. Sequence error refers to mismatches introduced during PCR or sequencing, which Bartender attempts to correct. In the ideal case, Bartender would cluster reads while correcting for all sequencing errors and introducing no additional error.

To evaluate the Bartender accuracy, a large simulated dataset with a defined sequence error rate, or a real dataset (Levy et al., 2015), is randomly partitioned into *n* equal subsets to mimic the sampling process. Bartender is then applied to each partition. For simulated data, Bartender results are compared to the true cluster centers known from the simulation; For real data, these results are compared to clusters that were generated by running Bartender on the entire data set prior to partitioning.

Since each read is sampled individually, for a particularly cluster, its size in each partition follows a Binomial distribution (equation 8).

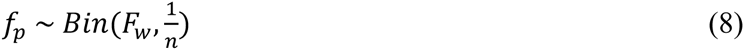

where *F_w_* is the size of the corresponding cluster in whole dataset.

So the expected cluster size in one partition is given by

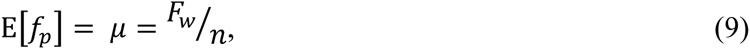

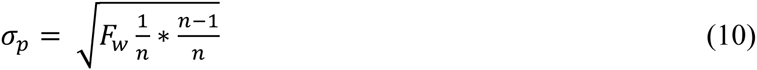

and the standard deviation in one partition is given by

The coefficient of variation (CV) (equation 11) and its estimator (equation 12) are used to gauge the error effects.

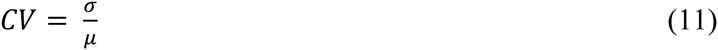

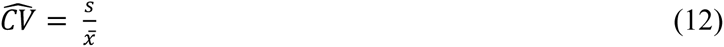

where 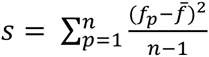, 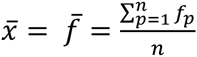 and *f_p_* is the observed frequency of each cluster in partition *p*.

### Sequence comparisons between BLAST and Bartender on a real data

We use ~130M reads of a 26mer barcode library (Levy et al., 2015) to compare BLAST and Bartender performance on real data. Here, the barcode library was generated from a random primer and, in contrast with simulated data, some barcodes are by chance longer or shorter than the expected 26 bases. Bartender uses a comparison metric based on the Hamming distance that does not allow comparisons between barcodes of different lengths, and instead clusters barcodes of different lengths independently. In contrast, BLAST uses a comparison metric based on the Levenshtein distance that allows barcodes of different lengths to be compared directly. This difference complicates comparisons between Bartender and BLAST, because BLAST cluster centers (the consensus nucleotide at each position) can be influenced by “frame-shifts” where reads with an additional or missing base at one position causes a misalignment in many other positions. Therefore, to maximize the overlaps between Bartender and BLAST results, we redefine the BLAST cluster center as the most frequent unique read within each cluster. Comparisons between BLAST and Bartender clusters are then performed using this definition of the BLAST cluster center.

### Multiple time point mode

For time course bar-seq data, Bartender first clusters data at each time point separately, then orders these clustering results by time point for further processing (Figure 8). For a time point *t*, Bartender first joins clusters with the exact same center (sequence) to *t*-1. Then, any unjoined clusters in *t* are used a query to clusters in *t*-1 if their centers that are within one mismatch. If no match is found and the size of cluster in time *t* is reasonably large (larger than a user-defined threshold), it will be recorded as a true barcode for time point t. If only one match is found in *t*-1, these barcode clusters are joined through time. If more than one match is found in *t*-1, the cluster in time t is joined with the largest cluster found in time t-1. This process is performed iteratively for all time points. Since the size of each barcode cluster has been recorded for all time points, with “zeros” for clusters absent at any time point, the final result contains a list of clusters and their counts at each time point (lineages). The overall cluster quality and position weight matrix across all time points are also included in the final report. More details are presented in the Algorithms 3 and 4 in Supplement.

**Figure 8.**
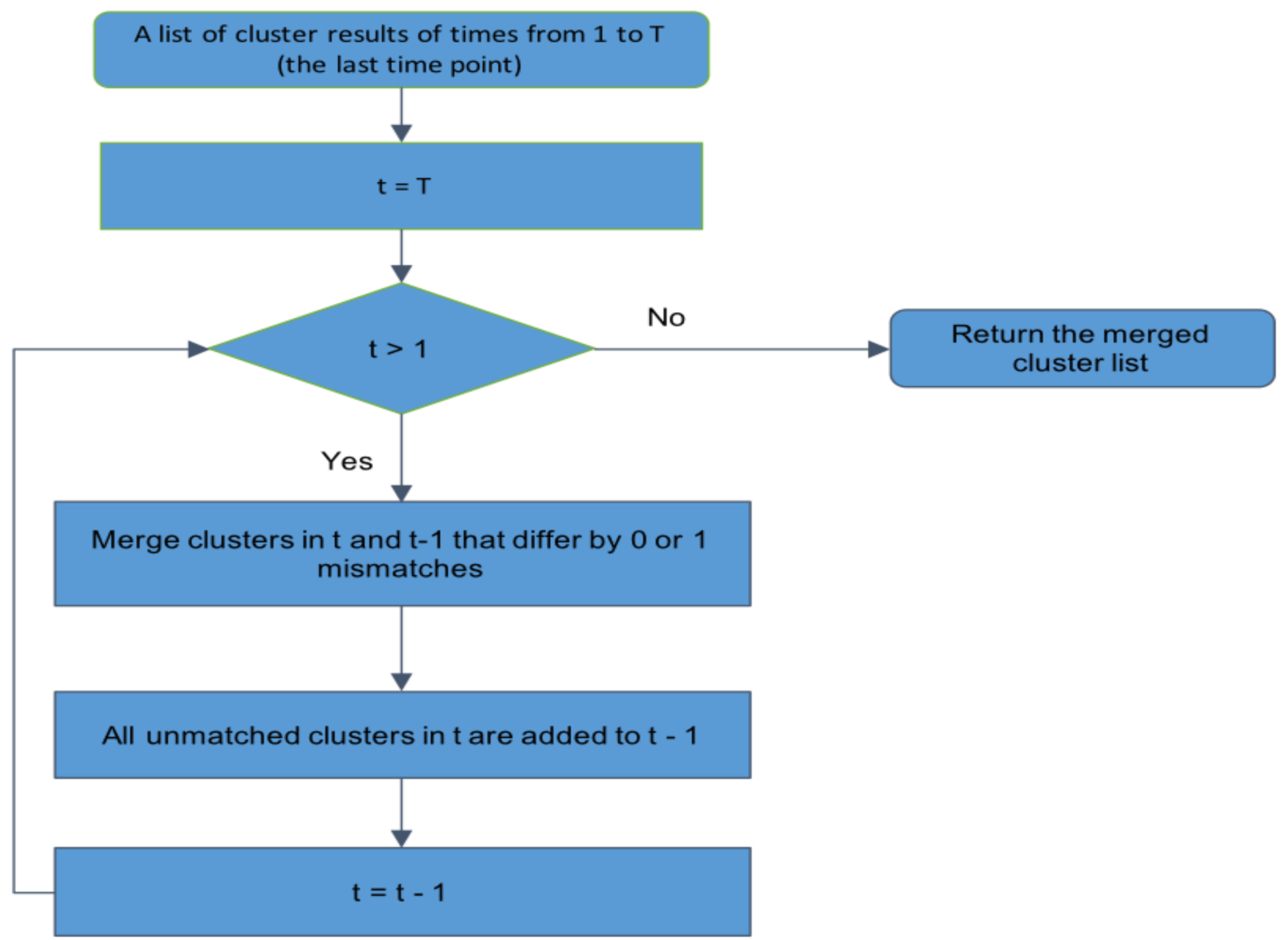
A flowchart of cluster merging across multiple time points. The merging strategy starts from a list of clustering results of each time point. Clusters from the latest available time point are merged to identical clusters from the previous time point. Clusters without an identical match merged with the most appropriate (see the main text) cluster within one mismatch. Other unmatched clusters will be kept only if its size is larger than the user specified size threshold.

## References

Altschul, Stephen F., Gish, Warren, Miller, Webb, Myers, Eugene W., & Lipman, David J. (1990). Basic local alignment search tool. Journal of Molecular Biology, 215(3), 403–410. 10.1016/s0022-2836(05)80360-2

Bassik, M. C., Lebbink, R. J., Churchman, L. S., Ingolia, N. T., Patena, W., LeProust, E. M.,… McManus, M. T. (2009). Rapid creation and quantitative monitoring of high coverage shRNA libraries. Nat Methods, 6(6), 443–445. 10.1038/nmeth.1330

Bhang, H. E., Ruddy,D. A., Krishnamurthy Radhakrishna, V., Caushi, J. X., Zhao, R., Hims, M. M.,… Stegmeier, F. (2015). Studying clonal dynamics in response to cancer therapy using high-complexity barcoding. Nat Med, 21(5), 440–448. 10.1038/nm.3841

Blundell, J. R., & Levy, S. F. (2014). Beyond genome sequencing: lineage tracking with barcodes to study the dynamics of evolution, infection, and cancer. Genomics, 104(6 Pt A), 417–430. 10.1016/j.ygeno.2014.09.005

Brutinel, E. D., & Gralnick, J. A. (2012). Anomalies of the anaerobic tricarboxylic acid cycle in Shewanella oneidensis revealed by Tn-seq. Mol Microbiol, 86(2), 273–283. 10.1111/j.1365-2958.2012.08196.x

Carette, J. E., Raaben, M., Wong, A. C., Herbert, A. S., Obernosterer, G., Mulherkar, N.,… Brummelkamp, T. R. (2011). Ebola virus entry requires the cholesterol transporter Niemann-Pick C1. Nature, 477(7364), 340–343. 10.1038/nature10348

Gawronski, J. D., Wong, S. M., Giannoukos, G., Ward, D. V., & Akerley, B. J. (2009). Tracking insertion mutants within libraries by deep sequencing and a genome-wide screen for Haemophilus genes required in the lung. Proc Natl Acad Sci U S A, 106(38), 1642216427. 10.1073/pnas.0906627106

Giaever, G., Chu, A. M., Ni, L., Connelly, C., Riles, L., Veronneau, S.,… Johnston, M. (2002). Functional profiling of the Saccharomyces cerevisiae genome. Nature, 418(6896), 387–391. 10.1038/nature00935

Gibney, P. A., Lu, C., Caudy, A. A., Hess, D. C., & Botstein, D. (2013). Yeast metabolic and signaling genes are required for heat-shock survival and have little overlap with the heat-induced genes. Proc Natl Acad Sci U S A, 110(46), E4393–4402. 10.1073/pnas.1318100110

Goodman, D. B., Church, G. M., & Kosuri, S. (2013). Causes and effects of N-terminal codon bias in bacterial genes. Science, 342(6157), 475–479. 10.1126/science.1241934

Goren, A., Ozsolak, F., Shoresh, N., Ku, M., Adli, M., Hart, C.,… Bernstein, B. E. (2010). Chromatin profiling by directly sequencing small quantities of immunoprecipitated DNA. Nat Methods, 7(1), 47–49. 10.1038/nmeth.1404

Gresham, D., Boer, V. M., Caudy, A., Ziv, N., Brandt, N. J., Storey, J. D., & Botstein, D. (2011). System-level analysis of genes and functions affecting survival during nutrient starvation in Saccharomyces cerevisiae. Genetics, 187(1), 299–317. 10.1534/genetics.110.120766

Gundry, M., & Vijg, J. (2012). Direct mutation analysis by high-throughput sequencing: from germline to low-abundant, somatic variants. Mutat Res, 729(1-2), 1–15. 10.1016/mrfmmm.2011.10.00110.1016/j.mrfmmm.2011.10.001

Hamming, R. W. (1950). Error Detecting and Error Correcting Codes. Bell System Technical Journal, 29(2), 147–160. 10.1002/j.1538-7305.1950.tb00463.x

Han, T. X., Xu, X. Y., Zhang, M. J., Peng, X., & Du, L. L. (2010). Global fitness profiling of fission yeast deletion strains by barcode sequencing. Genome Biol, 11(6), R60. 10.1186/gb-2010-11-6-r60

Hobbs, E. C., Astarita, J. L., & Storz, G. (2010). Small RNAs and small proteins involved in resistance to cell envelope stress and acid shock in Escherichia coli: analysis of a bar-coded mutant collection. J Bacteriol, 192(1), 59–67. 10.1128/JB.00873-09

Kivioja, T., Vaharautio, A., Karlsson, K., Bonke, M., Enge, M., Linnarsson, S., & Taipale, J. (2012). Counting absolute numbers of molecules using unique molecular identifiers. Nat Methods, 9(1), 72–74. 10.1038/nmeth.1778

Kosuri, S., Goodman, D. B., Cambray, G., Mutalik, V. K., Gao, Y., Arkin, A. P.,… Church, G. M. (2013). Composability of regulatory sequences controlling transcription and translation in Escherichia coli. Proc Natl Acad Sci U S A, 110(34), 14024–14029. 10.1073/pnas.1301301110

LeProust, E. M., Peck, B. J., Spirin, K., McCuen, H. B., Moore, B., Namsaraev, E., & Caruthers, M. H. (2010). Synthesis of high-quality libraries of long (150mer) oligonucleotides by a novel depurination controlled process. Nucleic Acids Res, 38(8), 2522–2540. 10.1093/nar/gkq163

Levenshtein, V. I. (1966). Binary Codes Capable of Correcting Deletions, Insertions and Reversals. Soviet Physics Doklady, 10.

Levy, S. F., Blundell, J. R., Venkataram, S., Petrov, D. A., Fisher, D. S., & Sherlock, G. (2015). Quantitative evolutionary dynamics using high-resolution lineage tracking. Nature, 519(7542), 181–186. 10.1038/nature14279

Lu, R., Neff, N. F., Quake, S. R., & Weissman, I. L. (2011). Tracking single hematopoietic stem cells in vivo using high-throughput sequencing in conjunction with viral genetic barcoding. Nat Biotechnol, 29(10), 928–933. 10.1038/nbt.1977

McKenna, A., Findlay, G. M., Gagnon, J. A., Horwitz, M. S., Schier, A. F., & Shendure, J. (2016). Whole organism lineage tracing by combinatorial and cumulative genome editing. Science. 10.1126/science.aaf7907

Meyerhans, Andreas, Vartanian, Jean-Pierre, & Wain-Hobson, Simon. (1990). DNA recombination during PCR. Nucleic Acids Research, 18(7), 1687–1691. 10.1093/nar/18.7.1687

Nguyen, L. V., Pellacani, D., Lefort, S., Kannan, N., Osako, T., Makarem, M.,… Eaves, C. J. (2015). Barcoding reveals complex clonal dynamics of de novo transformed human mammary cells. Nature, 528(7581), 267–271. 10.1038/nature15742

Noble, M., Treadwell, J. R., Tregear, S. J., Coates, V. H., Wiffen, P. J., Akafomo, C., & Schoelles, K. M. (2010). Long-term opioid management for chronic noncancer pain. Cochrane Database Syst Rev(1), CD006605. 10.1002/14651858.CD006605.pub2

Schlabach, M. R., Luo, J., Solimini, N. L., Hu, G., Xu, Q., Li, M. Z.,… Elledge, S. J. (2008). Cancer proliferation gene discovery through functional genomics. Science, 319(5863), 620–624. 10.1126/science.1149200

Schmitt, M. W., Kennedy, S. R., Salk, J. J., Fox, E. J., Hiatt, J. B., & Loeb, L. A. (2012). Detection of ultra-rare mutations by next-generation sequencing. Proc Natl Acad Sci U S A, 109(36), 14508–14513. 10.1073/pnas.1208715109

Schwarzmuller, T., Ma, B., Hiller, E., Istel, F., Tscherner, M., Brunke, S.,… Kuchler, K. (2014). Systematic phenotyping of a large-scale Candida glabrata deletion collection reveals novel antifungal tolerance genes. PLoS Pathog, 10(6), e1004211. 10.1371/journal.ppat.1004211

Silva, S. S., Rowntree, R. K., Mekhoubad, S., & Lee, J. T. (2008). X-chromosome inactivation and epigenetic fluidity in human embryonic stem cells. Proc Natl Acad Sci U S A, 105(12), 4820–4825. 10.1073/pnas.0712136105

Sims, D., Mendes-Pereira, A. M., Frankum, J., Burgess, D., Cerone, M. A., Lombardelli, C.,… Lord, C. J. (2011). High-throughput RNA interference screening using pooled shRNA libraries and next generation sequencing. Genome Biol, 12(10), R104. 10.1186/gb-2011-12-10-r104

Smith, G. J., Vijaykrishna, D., Bahl, J., Lycett, S. J., Worobey, M., Pybus, O. G.,… Rambaut, A. (2009). Origins and evolutionary genomics of the 2009 swine-origin H1N1 influenza A epidemic. Nature, 459(7250), 1122–1125. 10.1038/nature08182

van Opijnen, T., Bodi, K. L., & Camilli, A. (2009). Tn-seq: high-throughput parallel sequencing for fitness and genetic interaction studies in microorganisms. Nat Methods, 6(10), 767–772. 10.1038/nmeth.1377

Wang, T., Wei, J. J., Sabatini, D. M., & Lander, E. S. (2014). Genetic screens in human cells using the CRISPR-Cas9 system. Science, 343(6166), 80–84. 10.1126/science.1246981

Winzeler, E. A., Shoemaker, D. D., Astromoff, A., Liang, H., Anderson, K., Andre, B.,… Davis, R. W. (1999). Functional characterization of the S. cerevisiae genome by gene deletion and parallel analysis. Science, 285(5429), 901–906. 10.1126/science.285.5429.901

Wong, A. S., Choi, G. C., Cheng, A. A., Purcell, O., & Lu, T. K. (2015). Massively parallel high-order combinatorial genetics in human cells. Nat Biotechnol, 33(9), 952–961. 10.1038/nbt.3326

